# Closing Target Trimming: a Perl Package for Discovering Hidden Superfamily Loci in Genomes

**DOI:** 10.1101/490490

**Authors:** Zhihua Hua, Matthew J. Early

**Author notes:** Corresponding author: Zhihua Hua, Department of Environmental and Plant Biology, Ohio University, Athens, Ohio 45701 USA.

## Abstract

The contemporary capacity of genome sequence analysis significantly lags behind the rapidly evolving sequencing technologies. Retrieving biological meaningful information from an ever-increasing amount of genome data would be significantly beneficial for functional genomic studies. For example, the duplication, organization, evolution, and function of superfamily genes are arguably important in many aspects of life. However, the incompleteness of annotations in many sequenced genomes often results in biased conclusions in comparative genomic studies of superfamilies. Here, we present a Perl software, called Closing Target Trimming, for automatically identifying most, if not all, members of a gene family in any sequenced genomes. Our test data on the *F-box* gene superfamily showed 78.2 and 79% gene finding accuracies in two well annotated plant genomes, *Arabidopsis thaliana* and rice, respectively. This annotation performance is clearly higher than the best *ab initio* methods that are currently available. To further demonstrate the effectiveness of this program, we ran it through 18 plant genomes and five non-plant genomes to compare the expansion of the *F-box* and the *BTB* superfamilies. The program discovered that on average 12.7 and 9.3% of the total *F-box* and *BTB* members, respectively, are new loci in plant genomes while it only found a small number of new members in vertebrate genomes. Therefore, different evolutionary and regulatory mechanisms of cullin-RING ubiquitin ligases may be present in the plant and the animal kingdoms. Further studies may shed light on new discoveries in the ubiquitin-26S proteasome system-mediated regulatory pathways in eukaryotic organisms. With a detailed compiling instruction and a simple running operation, we expect that this software will assist many biological scientists with little programming experience to smoothly obtain a comprehensive dataset of a gene superfamily in any sequenced eukaryotic genomes.

## Introduction

Genome growth and innovation is largely attributed to the expansion of gene superfamilies [1-3]. Through both whole-genome duplication (WGD) and small-scale duplication (SSD) events, members of a gene family can increase dramatically [4-6]. Newly synthesized members in a family have been commonly recognized to experience either pseudogenization or fixation in a genome through four processes: 1) conservation if gene dosage is beneficial [1], 2) neofunctionalization if a novel function is acquired in one copy [1], 3) subfunctionalization if daughter copies split the function of the ancestral copy [7], or 4) specialization if all daughter copies perform new functions differing from their ancient copy [8]. Previous studies have suggested that dosage-balance constraints guarded duplicated copies against loss by determining the stoichiometry of duplicated products [9, 10]. Interestingly, families involving different cellular functions have various birth over death ratios. Transcription factors tend to have a much higher fixation rate than others, such as G-protein-coupled chemosensory receptors for finding food, odorants and pheromones and the immunoglobulins involving in the primary immune defense in vertebrates, and the nucleotide-binding site-leucine-rich repeat (NBS-LRR) receptors whose functions are largely found in defending pathogens in plants [11-16]. The significant contribution in genome evolution and the striking variance of evolutionary processes of members within and between families place the studies of gene duplication an important role in decoding the function of genomes. To reach this, precisely finding the total members of a gene family in a genome would be the first effort.

Since the era of draft sequencing of human genome, finding genes in massive genomic DNA sequences has drawn significant attention in understanding the structure and function of genomes [17]. Early in this century, many computational methods were developed in order to predict genes in a piece of DNA or in an entire genome precisely. Overall, these methods can be divided into two categories: sequence similarity-based extrinsic methods and machine-learning curated intrinsic approaches. The former employs known query sequences to align a target sequence to discover the structure of an unknown locus while the latter utilizes dynamic programming to identify genes by locating the gene elements (e.g., specific signals in the 5’-or 3’-ends of genes) [18]. Due to the extreme variability of gene elements, the intrinsic (*ab initio*) methods often yield more false predictions than the extrinsic evidence-based prediction [18]. For example, prior *ab initio* methods predicted *Caenorhabditis elegans* and *Homo sapiens* genes with an accuracy rate of 50% and 20%, respectively [19, 20]. Certainly, the accuracy of both types of gene annotation methods has been significantly increased since both the scale and the detail of known genes are now dramatically expanded and improved. Recently developed next generation mRNA sequencing-based methods have increased the accuracy of *de novo* genome annotations to an average of 57.6% on *C. elegans* genes [21, 22]. However, the human ENCODE genome annotation assessment project, EGASP, suggested that the mRNA and protein sequences-based annotation programs still yielded the most accurate prediction of gene models [20].

With the booming of next generation sequencing technologies [23], large-scale genome sequencing has become routine in many individual laboratories while genome annotation and analysis remains challenging. Considering the importance of gene superfamily studies and the contemporary large accumulation of accomplished genome projects, we developed a Perl package, named Closing Target Trimming (CTT), for superfamily annotation. This package incorporates the well-established sequence similarity-based GeneWise annotation tool [24] to automatically identify and re annotate most, if not all, members of a gene superfamily in a genome, thus benefiting comparative genomic studies.

The algorithm of CTT was first developed during the course of studying the phylogenetics of the *F-box* gene superfamily in plants [5]. When we retrieved the *F-box* gene members in prior annotations from 18 sequenced plant genomes, including both seedless and seed plants, we noticed that there were many novel *F-box* putative loci left unannotated in genomes. Those loci are either lacking clear gene signals that allow them to be discovered *ab initio* or too close with one another to be uncovered by GeneWise annotation, since the latter only finds the best homologue of a known query protein sequence [5, 24]. Recently, we have also employed this algorithm to study the duplication mechanism of ubiquitin and ubiquitin-like protein (in whole called ubiquiton) family in 50 sequenced plant genomes [6]. CTT has been demonstrated as an effective tool to retrieve most, if not all, members of a gene superfamily in genomes we have analyzed. To make this program readily available for the biological science community, we organized and improved the scripts used in the previous two studies and compiled it into one integrated package that allows researchers with little programming experience to use.

## Materials and Methods

### Package development

The package is written in Perl (Version 5) and has been tested on CentOS 7 Linux operating system.

### Similarity-Based Gene Annotation

Based on our previous studies on CTT-based re-annotation of *F-box* and *ubiquiton* genes in plant genomes [5, 6], we designed to retrieve a 10,000 nucleotide (nt) genomic sequence sandwiching the start coordinate of a target domain for identifying a novel family locus. Since a non-plant gene may be longer than a plant gene on size, we also examined the effectiveness of CTT prediction by expanding the size of a putative locus to 100,000 nts. A putative novel locus is positioned by tBLANTn search [25] using the query of a collection of seed sequences, which represent the domain signature of a target family, against the genome sequence. Only if a new locus identified by CTT iteration search (see algorithm below) encompasses the tBLASTn-defined locus would the genomic sequence be further analyzed. To eliminate a potential false prediction due to the inaccuracy of one reference protein sequence, we applied at least the top two best homologous protein sequences (based on bit scores of BLASTx search [25]) to back search the genomic DNA sequence for predicting the transcript model of a putative novel locus using GeneWise [24]. If the locus is consistently predicted as a protein-coding gene by at least two reference sequences, the coding sequence identified with the highest GeneWise score was taken as the best prediction of this putative new locus. The resulting peptide sequence is further examined for the presence of a target family domain using HMMER search (http://hmmer.org) against the Pfam-A database (Version 32.0, https://pfam.xfam.org, E-value <1).

### CTT Iteration Search Algorithm

The genomic DNA sequence with 10,000 nts is likely to contain more than one locus. This is particularly common in many superfamily genes that were tandemly duplicated. This prevents the identification of a putative new locus by GeneWise because GeneWise only aligns a reference protein sequence to the best homologous region [24]. To expose this putative new locus defined by tBLASTn search, unrelated adjacent DNA sequence is iteratively trimmed and removed by CTT. First, the 10,000 nt genomic DNA sequence is used to identify a locus that best matches to a homologous protein sequence by GeneWise (GeneWise score ≥ 50). The DNA sequence of a predicted locus is trimmed and removed if it does not encompass the tBLASTn-defined region. The remaining DNA sequence is then iterated for a sequential BLASTx-GeneWise-trimming search until the tBLASTn-defined region is identified to be within the GeneWise predicted locus. Otherwise, the tBLASTn-defined region is not considered to be a new family locus.

### Data Resources

To test the effectiveness of this CTT package, a genome sequence file, a generic feature format 3 (GFF3) file, and a protein sequence file from 18 sequenced plant genomes and 5 sequenced non-plant genomes (Table S1) were retrieved from Phytozome (Version 12, https://phytozome.jgi.doe.gov) and Ensembl (Release 94, https://useast.ensembl.org), respectively. The seed files for two test superfamilies, the F-box (PF00646) and the BTB (PF00651) protein families, and the Pfam-A database were downloaded from Pfam (Version 32.0, https://pfam.xfam.org).

## Results

### Structure and Implementation

Currently, the package is built on CentOS 7 Linux operating system. The source code is permitted to use and distribute under the GNU General Public License Version 3.0. The package requires the Pfam-A database and six program dependencies, including BioPerl (https://bioperl.org), BLAST [25], HMMER (http://hmmer.org), GeneWise [24], PfamScan (https://pfam.xfam.org) and CD-HIT [26]. The users are strongly recommended to follow the steps described in a *README.md* file under the main directory /*ctt* to install the database and the dependencies on CentOS 7. We found that compiling BioPerl could be really problematic for researchers with little programming experience. The instructions provided in “*Install BioPerl*” in the *README.md* file can significantly reduce this hassle. After BioPerl and BioPerl-Run are installed, the “*make all*” function in *makefile* under the directory */ctt/dependencies* should help to install the Pfam-A database and the remaining program dependencies smoothly.

Before running the program, the users need to collect the information of target genomes and seed sequences for a family of interest as input files into the directories */ctt/species_databases* and */ctt/seeds*, respectively. Since we are interested in both known and unknown members of a family, a genome sequence file, a generic feature format 3 (GFF3) file, and a protein sequence file from a previous genome annotation are required to be deposited in the directory of */ctt/species_databases*. The family members from prior annotations are used as reference peptide (Ref_Pep) sequences to predict the gene structure of an unknown locus using GeneWise. Therefore, we recommend adding the same set of three files from well-annotated genomes to increase the prediction accuracy. The package is capable of working on multiple genomes at one time. Once the number of genomes is decided and all the required information is collected, a tab file, named “*organismal_genome_gff3_proteome_files.tab*”, needs to be prepared with each row containing the names of a genome sequence file, a GFF3 file, and a protein sequence file of one genome, each of which is separated with a tab space. The seed file of a family can be found and downloaded on Pfam (https://pfam.xfam.org) and should be placed in the directory */ctt/seeds*. All the input sequence files are in a FASTA format, which allows a BioPerl module, Bio::DB::Fasta, to parse the sequences. These files should also be converted to BLASTable databases in the same directory using the *makeblastdb* function as instructed in the *README.md* file or as described in the manual of BLAST [25].

The operation code of this program is formulated as one Perl command in the terminal under the directory of the package */ctt*. The users simply need to enter a code formatted as “*perl ctt.pl-seed family_seed_file.txt-f Pfam_family_id -superfamily simplified_family_id_you_named*” to start the program. The *ctt.pl* should automatically read the input files from */ctt/species_databases* and */ctt/seeds* to finish the annotation. There are in total seven steps, each of which is operated by an annotation module saved in the directory*/ctt/annotation_modules*. In accordance, there are 7 outputs that the program produces and saves stepwise in a final */ctt/ctt_output* directory (Table 1).

**Table 1.**
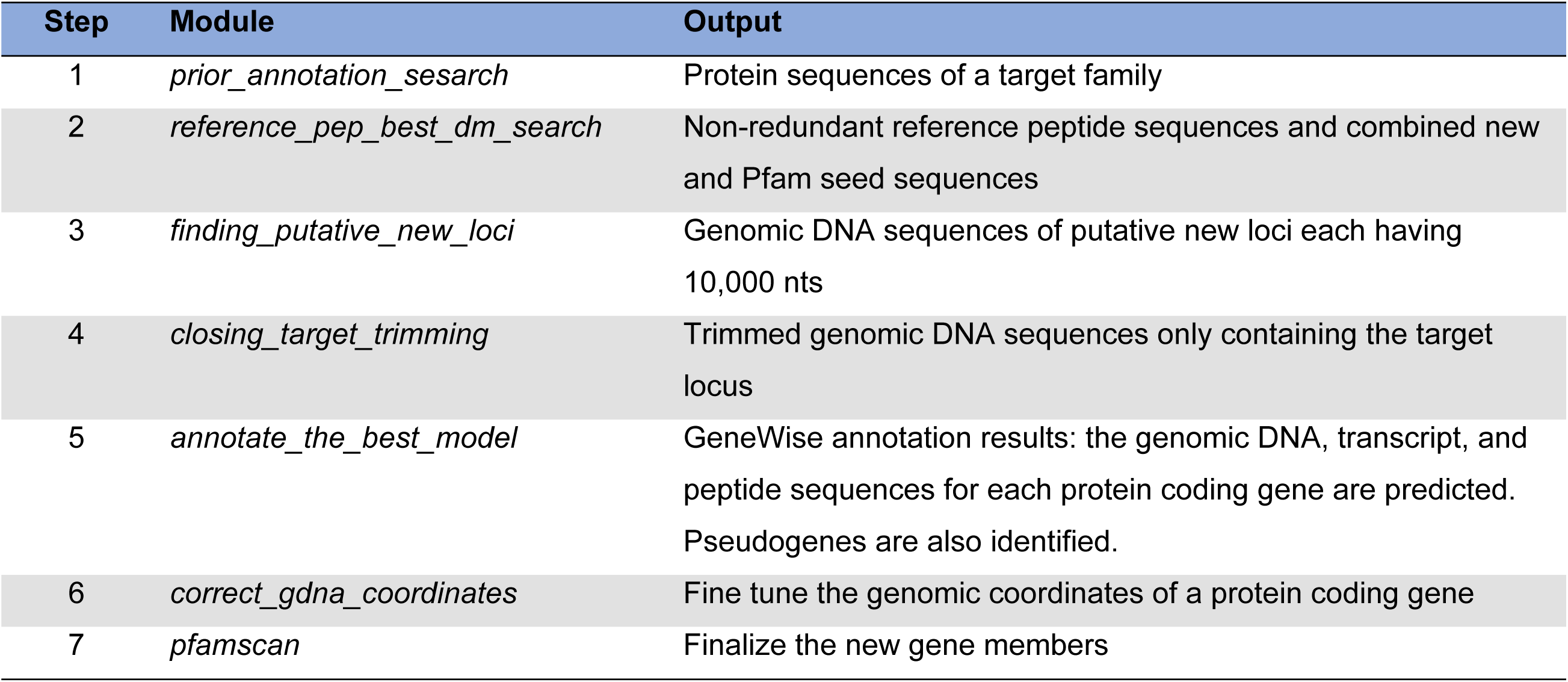
Annotation modules and outputs throughout the running process of the package.

In Step 1, *prior_annotation_search.pm* applies the seed sequences as query to BLASTp [25] a previously-annotated protein database. Pfam domains present in each hit protein sequence are obtained by *PfamScan.pl* search against the Pfam-A database that is automatically downloaded, processed, and saved under the directory */ctt/databases/pfam*. The Pfam domain search in *PfamScan.pl* is essentially done by HMMER3 (http://hmmer.org). The latter applies profile hidden Markov models (profile HMMs) to determine the presence of a sequence homologous to a Pfam domain, whose information is represented by a multiple sequence alignment and a hidden Markov model (HMM) in the Pfam-A database (https://pfam.xfam.org). In the current version of CTT, we apply 1e-5 and 1 as an E-value cutoff for the BLASTp and HMMER searches, respectively. Both values are artificially determined based on our previous studies that may assist to include most known members in both the *F-box* and the *ubiquiton* families [5, 6].

The module of Step 2, named *reference_pep_best_dm_search.pm*, is designed to organize the full-length protein sequences of family members predicted in Step 1 and retrieve the best family domain sequence in each protein sequence as a new seed sequence. Both the full-length sequences and the combined new and Pfam family seed sequences are subject to CD-HIT [26] to remove redundant ones. The retaining full-length and seed sequences are used as reference and seed sequences for later GeneWise annotation and tBLASTn search, respectively.

Step 3 applies the non-redundant seed sequences as queries to search a genome database using tBLASTn with an e-value cutoff ≤ 1e-5. Since the tBLASTn search generates a large number of hits, we considered two hits the same if the middle coordinate of one resides in the region of the other. In addition, we use AWK (comes with centOS) to compare the start and the end coordinates of one tBLASTn hit with those of the previously-annotated genes, whose information is retrieved from the GFF3 file. Since tBLASTn finds a region aligned with a family domain, the hit should reside in an annotated gene if it is part of the gene. Those hits are not considered as new loci and are removed from further analyses. This step is accomplished using a module named *finding_putative_new_loci.pm*.

Step 4 runs the CTT algorithm which is coded in a module called *closing_target_trimming.pm*. In order to gain a genomic region long enough to cover a putative new locus, we added a sequence of 5,000 nts on both sides of the start point of a retaining tBLASTn hit in Step 3. Based on our previous study on *F-box* genes, the longest *F-box* locus in plants is less than 10,000 nts [5]. A DNA sequence of 10,000 nts should be sufficient to contain a new locus in plants. For other organisms, the users can modify *finding_putative_new_loci.pm* in Step 3 to retrieve a longer region if needed. This 10,000 nt DNA sequence containing an unknown locus is used to BLASTx the Ref_Pep database obtained in Step 2. The best hit is then used as a protein template in GeneWise to predict the structure of a new gene. However, the best homology search algorithm may not let GeneWise to find the locus containing the target hit if there are multiple homologous sequences in this DNA fragment. To expose the target locus, this module applies up to six iterations of GeneWise annotation and trimming to shorten the genomic sequence until a hit-containing locus is discovered (Figure 1). The iteration stops if the GeneWise score is lower than 50 or no BLASTx hit is obtained.

**Figure 1.**
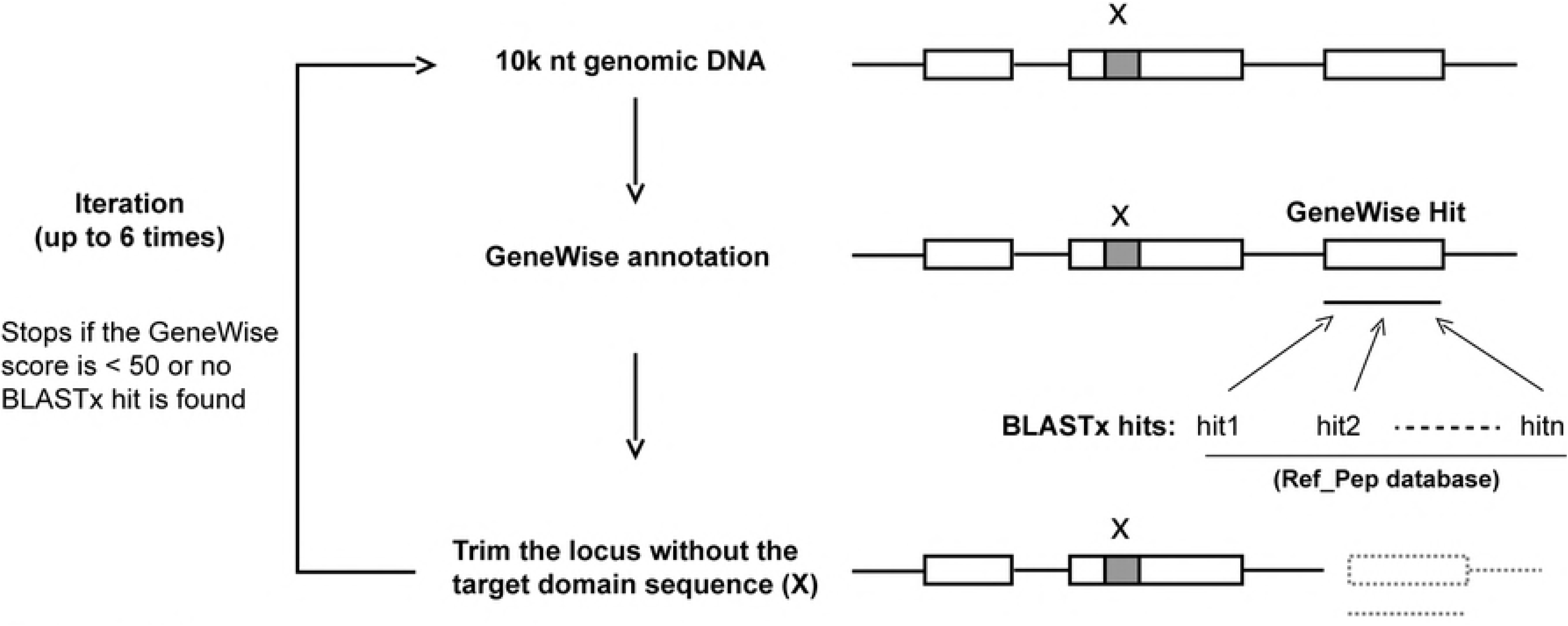
A schematic illustration of the CTT algorithm. The “x” indicates a genomic region that encodes a target family domain. The coordinates of “x” is defined by a tBLASTn search using the query of seed sequences of the target family. “Hit1, 2,…n” indicate the top hits from a BLASTx search using a 10,000 nt genomic DNA sequence containing “x” as query against protein sequences of previously-annotated family members. The process is iterated for up to six times to shorten the genomic sequence until a new locus containing “x” is discovered. The iteration stops if the GeneWise score is lower than 50 or no BLASTx hit is obtained.

To reduce the bias caused by a problematic reference sequence, Step 5 calls the module, *annotate_the_best_model.pm*, to utilize the protein sequences of the three best BLASTx hits as GeneWise templates to annotate the gene structure of an unknown target locus uncovered in Step 4. Only if one of the three gene models (peptide-coding gene, pseudogene without GeneWise transcript, or pseudogene containing early stop codons) is consistently predicted by at least two reference sequences would we consider it a good annotation. The transcript predicted with the highest GeneWise score is taken as the final prediction for a peptide-coding gene.

Step 6 retrieves the correct genomic coordinates of a putative new member and Step 7 confirms the presence of a family domain in the new annotated protein sequence (Table 1).

### Performance Test

We tested both sensitivity and specificity of CTT on gene finding using the *F-box* gene family as an example in two well annotated plant genomes, *Arabidopsis thaliana* and rice (*Oryza sativa*). First, we used *prior_annotation_search.pm* to identify the full set of *F-box* gene members in the previously predicted proteomes of Arabidopsis and rice. To assess the performance, we randomly selected 100 *F-box* members in each species and removed their annotations in the corresponding proteome. We then ran through CTT to examine 1) how many of these 100 *F-box* genes were rediscovered in each genome, and 2) how good their annotations are compared to the previous protein sequences. To increase both the sensitivity and specificity of CTT, we collected the annotation datasets from 18 plant genomes (S1 Table) as we did previously [5].

We defined that a test *F-box* locus was identified if the midpoint coordinate of the CTT-annotated locus is within the test locus. We found that 75 and 85 test *F-box* loci were rediscovered by CTT in Arabidopsis and rice genomes, respectively. We also found by CTT that 15 and 20 test *F-box* loci from Arabidopsis and rice, respectively, were predicted to have no transcript available or express a protein with premature stop codons, indicative of pseudogenes (Table 2). This result further confirmed our previous discoveries showing that a significant number of *F-box* loci are pseudogenized [5, 10].

**Table 2.**
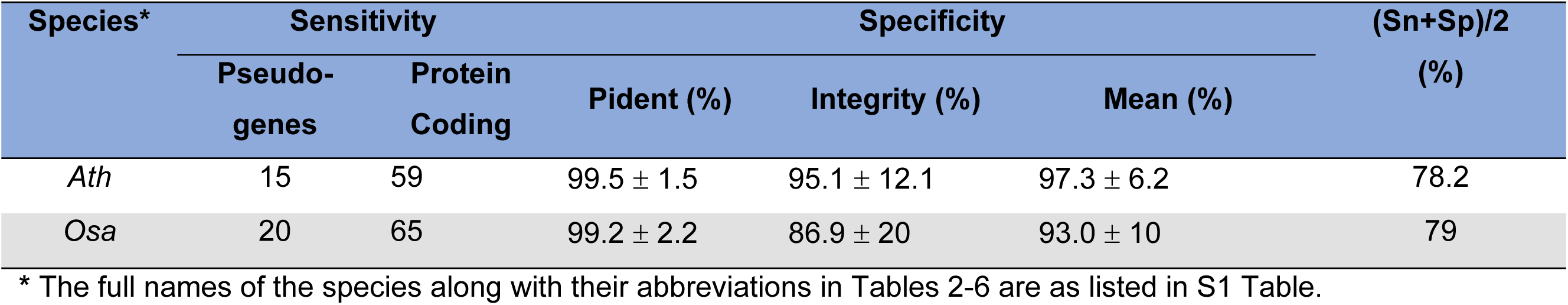
The sensitivity and specificity of CTT in finding plant *F-box* gene members.

| Species* | Sensitivity |  | Specificity |  |  | (Sn+Sp)/2 (%) |
| --- | --- | --- | --- | --- | --- | --- |
|  | Pseudo-genes | Protein Coding | Pident (%) | Integrity (%) | Mean (%) |  |
| *Ath* | 15 | 59 | 99.5 ± 1.5 | 95.1 ± 12.1 | 97.3 ± 6.2 | 78.2 |
| *Osa* | 20 | 65 | 99.2 ± 2.2 | 86.9 ± 20 | 93.0 ± 10 | 79 |

\* The full names of the species along with their abbreviations in Tables 2-6 are as listed in S1 Table.

To test the specificity of CTT, we compared both sequence identity and integrity of the predicted protein sequences with previous annotations using BLASTp analysis. Between the two genomes compared, CTT seems to perform slightly better in *Arabidopsis* to predict the full length of a test *F-box* locus (Table 2, *p* = 0.005, Student’s *t*-test). This is likely due to a close relationship between a reference genome, *A. lyrata*, and *A. thaliana*. However, for all the protein sequences predicted, CTT yielded >99% sequence identities with the original predictions, which are consistent with the outstanding performance of GeneWise annotation if a good reference sequence is available [24]. After averaging both specificity and sensitivity, CTT obtained 78.2 and 79% accuracy in finding a new *F-box* locus in Arabidopsis and rice, respectively (Table 2).

### Size Comparison of the *F-box* and the *BTB* Superfamilies in Plants

We then applied CTT to search the members of two gene superfamilies in 18 selected plant genomes. In addition to the *F-box* gene family, we selected the *bric-a-brac/tramtrack/broad complex* (*BTB*) family for comparison. The criteria of selecting these two families are 1) that both encode a substrate receptor in a cullin-RING E3 ubiquitin ligase complex important for regulating a vast number of metabolic and signaling pathways in all eukaryotic organisms [27], and 2) that both families are largely expanded in plant genomes [5, 28].

Consistent with our phylogenetic studies on the *F-box* gene superfamily, we obtained a similar number of *F-box* genes in all 18 plant genomes (*ρ* = 0.96, p-value = 6.7e-06, Spearman’s correlation test) (Figure 2A, S1-S4 Files). The slight difference in the number of *F-box* genes found in the two studies may have resulted from 1) the upgrade of genome and F-box seed sequences, and 2) the improvement of CTT in this package. For example, in this package, we incorporated GFF3 files to precisely locate known loci in a prior annotation (Step 3, *finding_putative_new_loci.pm*, Table 1). However, our previous study used BLAT search [29] to locate an annotated gene, which may result in inaccurate genomic coordinates [5].

**Figure 2.**
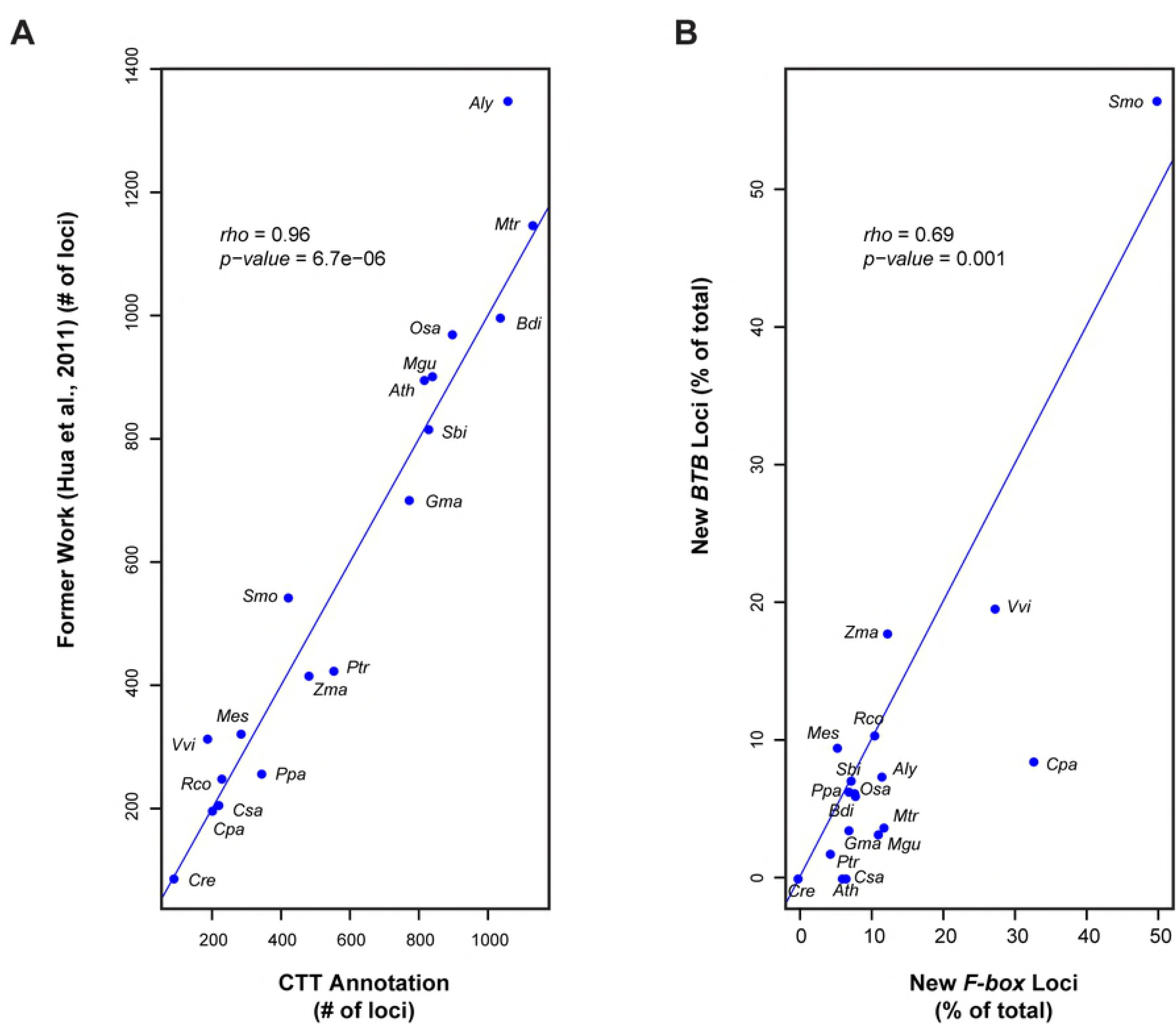
Performance test on the CTT program. (A) Annotation comparison of the *F-box* gene superfamily in 18 plant genomes between a previous work [5] and the output of CTT automatic annotation in this work. (B) Number correlation of new *F-box* and new *BTB* loci discovered in 18 plant genomes. The blue line indicates equal x axis and y axis values. The full names of the species along with their abbreviations are as listed in S1 Table.

After CTT annotation, we discovered that, on average, 12.7 and 9.3% of the total members in the *F-box* and *BTB* families, respectively, are new loci (Tables 3 and 4, S5-S8 Files). Although a slightly higher proportion of *F-box* genes were discovered than that of *BTB* genes (*p* < 0.05, Student’s *t*-test), the percentiles of new members in these two families are significantly correlated (*ρ* = 0.69, p-value = 0.001, Spearman’s correlation test) (Figure 2B). Therefore, there are various annotation qualities in different sequenced genomes, further highlighting the effectiveness of this package in helping identify most, if not all, superfamily members in a genome for comparative genomic studies.

**Table 3.** CTT annotation of *F-box* genes in 18 plant genomes.

| Species | Prior Annotation | CTT Annotation |  | Total | New Finding<br>(% of total) |
| --- | --- | --- | --- | --- | --- |
|  |  | Pseudogenes | Protein Coding<br>Genes |  |  |
| *Aly* | 940 | 65 | 59 | 1064 | 11.7 |
| *Ath* | 771 | 34 | 17 | 822 | 6.2 |
| *Bdi* | 960 | 43 | 39 | 1042 | 7.9 |
| *Cpa* | 139 | 53 | 15 | 207 | 32.9 |
| *Cre* | 95 | 0 | 0 | 95 | 0.0 |
| *Csa* | 210 | 3 | 12 | 225 | 6.7 |
| *Gma* | 723 | 40 | 15 | 778 | 7.1 |
| *Mes* | 274 | 5 | 11 | 290 | 5.5 |
| *Mgu* | 750 | 49 | 46 | 845 | 11.2 |
| *Mtr* | 1000 | 85 | 51 | 1136 | 12.0 |
| *Osa* | 831 | 35 | 37 | 903 | 8.0 |
| *Ppa* | 325 | 6 | 19 | 350 | 7.1 |
| *Ptr* | 534 | 18 | 7 | 559 | 4.5 |
| *Rco* | 209 | 12 | 13 | 234 | 10.7 |
| *Sbi* | 772 | 35 | 27 | 834 | 7.4 |
| *Smo* | 213 | 80 | 134 | 427 | 50.1 |
| *Vvi* | 140 | 15 | 38 | 193 | 27.5 |
| *Zma* | 426 | 38 | 23 | 487 | 12.5 |

**Table 4.** CTT annotation of *BTB* genes in 18 plant genomes.

| Species | Prior Annotation | CTT Annotation |  | Total | New Finding<br>(% of total) |
| --- | --- | --- | --- | --- | --- |
|  |  | Pseudogenes | Protein Coding<br>Genes |  |  |
| *Aly* | 75 | 2 | 4 | 81 | 7.4 |
| *Ath* | 86 | 0 | 0 | 86 | 0.0 |
| *Bdi* | 197 | 11 | 2 | 210 | 6.2 |
| *Cpa* | 43 | 2 | 2 | 47 | 8.5 |
| *Cre* | 70 | 0 | 0 | 70 | 0.0 |
| *Csa* | 55 | 0 | 0 | 55 | 0.0 |
| *Gma* | 194 | 4 | 3 | 201 | 3.5 |
| *Mes* | 67 | 3 | 4 | 74 | 9.5 |
| *Mgu* | 60 | 0 | 2 | 62 | 3.2 |
| *Mtr* | 105 | 2 | 2 | 109 | 3.7 |
| *Osa* | 173 | 7 | 4 | 184 | 6.0 |
| *Ppa* | 89 | 1 | 5 | 95 | 6.3 |
| *Ptr* | 163 | 2 | 1 | 166 | 1.8 |
| *Rco* | 43 | 2 | 3 | 48 | 10.4 |
| *Sbi* | 171 | 9 | 4 | 184 | 7.1 |
| *Smo* | 30 | 2 | 37 | 69 | 56.5 |
| *Vvi* | 45 | 7 | 4 | 56 | 19.6 |
| *Zma* | 125 | 20 | 7 | 152 | 17.8 |

### CTT Annotation of the *F-box* and the *BTB* Superfamilies in Non-plant Genomes

Since the CTT algorithm was first developed to study the *F-box* gene superfamily in plants [5], we questioned whether this newly designed CTT package was also able to annotate new family members in non-plant genomes. We selected the genomes of human and its two-close relatives, the chimpanzee and mouse, as well as nematode and budding yeast as a test dataset (S1 Table). For a strict comparison, we applied CTT to annotate the *F-box* and the *BTB* families in these five genomes.

Unlike the large and unstable expansion of the *F-box* superfamily in plants (Table 3 and [5]), we found that its size in three closely related vertebrates is fairly small and stable (Table 5, S9-S12 Files), suggesting that plants and vertebrates may have adapted a different mechanism in regulating the evolution and the function of the F-box-mediated protein ubiquitylation system. Consistent with the broad post-translational regulatory function of the ubiquitin-26S proteasome system (UPS), CTT annotation discovered that the three vertebrates have a larger BTB family than many flowering plant species (Table 6, S13-S16 Files). In addition, the expansion of this family in these genomes is like the *F-box* superfamily showing little size variation. More interestingly, only a small proportion of members from both families were newly discovered by CTT in the chimpanzee and worm genomes, and none were discovered in the other three non-plant genomes (Tables 5 and 6).

**Table 5.**
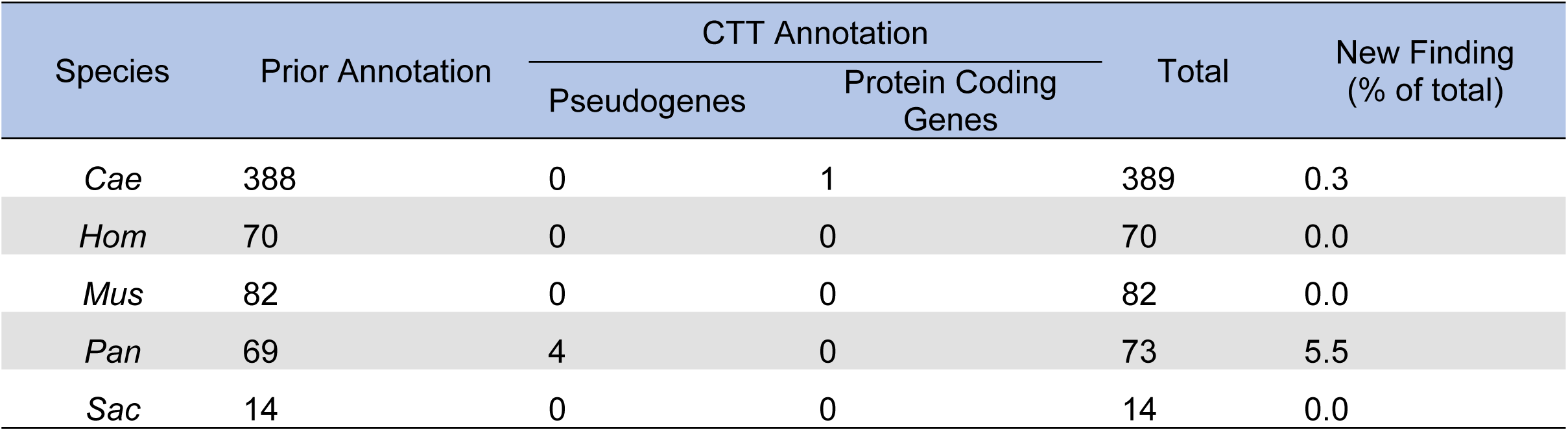
CTT annotation of *F-box* genes in 5 non-plant genomes.

**Table 6.** CTT annotation of *BTB* genes in 5 non-plant genomes.

| Species | Prior Annotation | CTT Annotation |  | Total | New Finding<br>(% of total) |
| --- | --- | --- | --- | --- | --- |
|  |  | Pseudogenes | Protein Coding<br>Genes |  |  |
| *Cae* | 126 | 0 | 0 | 126 | 0.0 |
| *Hom* | 151 | 0 | 0 | 151 | 0.0 |
| *Mus* | 183 | 2 | 0 | 185 | 1.1 |
| *Pan* | 155 | 6 | 4 | 165 | 6.1 |
| *Sac* | 1 | 0 | 0 | 1 | 0.0 |

We initially thought that the low number of new loci detected by CTT in these five non-plant genomes might be due to a larger genomic size of vertebrate genes than that of plant genes. We revised the *finding_putative_new_loci.pm* module in Step 3 to retrieve a genomic DNA sequence of 100,000 nts instead of 10,000 nts for downstream CTT annotation. This modification did not make an improvement and missed the only new locus discovered in *C. elegans* by the original approach. Therefore, we think a genomic DNA sequence of 10,000 nts would be a good size for searching the presence of an *F-box* or a *BTB* gene.

## Discussion

The invention of next-generation sequencing technologies revolutionized genomic studies by dramatically reducing the cost and increasing the speed in various genome sequencing projects. The recently-emerging third generation (or long-read) sequencing technologies would further this revolution by sequencing genomes in a much more affordable, accurate, and effective manner [23]. The contemporary genome sequencing technologies have already made it possible to sequence the genomes of every organism on the earth, such as the Earth BioGenome Project (EBP) initiated this year [30]. However, a sequencing machine only produces a lengthy genome book composed of 4 or 5 letters, i.e., A, T, C, G, or N. To understand the organization, evolution, functions, and interactions of numerous genomes, post genome sequencing data analysis is much more challenging than ever before. For example. many biological laboratories are not yet able to take full advantage of the power of whole-genome sequencing because of the lack of the bioinformatic tools and/or skills.

While many state-of-the-art bioinformatic tools have been developed to annotate genomes u, our studies have shown that superfamily members were often missing in various accomplished genome projects [5, 6] (Tables 3 and 4). Studying gene duplications of a family across genomes can shed light on the evolutionary and functional mechanisms of intracellular regulatory pathways [4, 6, 10]. Yet, a biased conclusion could be made if a comprehensive understanding of the members of a gene family in genomes is not achieved. For example, it was concluded that herbaceous annual plants have a larger *F-box* gene family than woody perennial plants from a study of few plant genomes [31]. However, our later CTT-based comprehensive re-annotation of the *F-box* genes in a broad range of plant genomes did not fully support this conclusion [5]. Our previous data and the data in this work clearly show that the size of the *F-box* superfamily is not significantly correlated with the growth behavior of a plant species, indicative of a genomic drift evolutionary mechanism ([5] and Table 3). The *F-box* family in herbaceous plants, papaya (*Carica papaya*), cucumber (*Cucumis sativus*), and *Castor bean* (*Ricinus communis*), is smaller than that in the woody plants *Populus trichocarpa* and *Medicago truncatula* (Table 3).

The effectiveness of CTT in annotating the *F-box* and the *BTB* superfamilies in both plants and non-plant organisms suggests a broad application of this software. The availability of this CTT package will benefit both evolutionary and functional genomic studies of superfamily genes in eukaryotic organisms. Our test data also suggested that different evolutionary mechanisms might have contributed differently to the expansion of the *F-box* and the *BTB* superfamilies in the plant and the animal kingdoms. It would be worth further comparing the duplication mechanisms of the entire ubiquitin-26S proteasome system in these two kingdoms in the future.

### Package availability

The full CTT package is available in a repository hosted by GitHub at https://github.com/hua-lab/ctt.

## Supporting information

**S1 Table. List of 18 plant genomes and 5 non-plant genomes used in this work.**

**S1 File. Protein sequences of the *F-box* genes identified by CTT in prior annotations of 18 plant species.**

**S2 File. Protein sequences of novel *F-box* genes reannotated by CTT in 18 plant species. S3 File. Transcript sequences of novel *F-box* genes reannotated by CTT in 18 plant species.**

**S4 File. Genomic DNA sequences of novel *F-box* genes reannotated by CTT in 18 plant species.**

**S5 File. Protein sequences of the *BTB* genes identified by CTT in prior annotations of 18 plant species.**

**S6 File. Protein sequences of novel *BTB* genes reannotated by CTT in 18 plant species. S7 File. Transcript sequences of novel *BTB* genes reannotated by CTT in 18 plant species.**

**S8 File. Genomic DNA sequences of novel *BTB* genes reannotated by CTT in 18 plant species.**

**S9 File. Protein sequences of the *F-box* genes identified by CTT in prior annotations of five non-plant species.**

**S10 File. Protein sequences of novel *F-box* genes reannotated by CTT in five non-plant species.**

**S11 File. Transcript sequences of novel *F-box* genes reannotated by CTT in five non-plant species.**

**S12 File. Genomic DNA sequences of novel *F-box* genes reannotated by CTT in five non-plant species.**

**S13 File. Protein sequences of the *BTB* genes identified by CTT in prior annotations of five non-plant species.**

**S14 File. Protein sequences of novel *BTB* genes reannotated by CTT in five non-plant species.**

**S15 File. Transcript sequences of novel *BTB* genes reannotated by CTT in five non-plant species.**

**S16 File. Genomic DNA sequences of novel *BTB* genes reannotated by CTT in five non-plant species.**

## Acknowledgments

We thank each genome-sequencing project for providing the annotation and genome sequences and the Pfam database for its data resources. We also thank numerous researchers for providing the free open source codes in the five dependencies adopted in this package. This work was supported by a National Science Foundation CAREER award (MCB-1750361 to ZH) and Ohio University Program to Aid Career Exploration (PACE) fund (ZH9299110 to ZH).

## Author Contributions

**Conceptualization**: Zhihua Hua

**Data Curation**: Zhihua Hua

**Formal Analysis**: Zhihua Hua and Matthew Early

**Funding Acquisition**: Zhihua Hua

**Investigation**: Zhihua Hua

**Methodology**: Zhihua Hua and Matthew Early

**Software**: Zhihua Hua

**Supervision**: Zhihua Hua

**Visualization**: Zhihua Hua

**Writing – original draft**: Zhihua Hua

**Writing – review & editing**: Zhihua Hua

## References

1. Ohno S. Evolution by Gene Duplication. Springer-Verlag, New York. 1970.

2. Li. W-H. Evolution of duplicate genes and pseudogenes. In: Evolution of Genes and Proteins (M Nei, and RK Koehn eds). 1983:Sinauer Associates, Sunderland, MA. pp 14–37.

3. Panchy N, Lehti-Shiu M, Shiu SH. Evolution of Gene Duplication in Plants. Plant Physiol. 2016;171(4):2294–316. Epub 2016/06/12. doi: 10.1104/pp.16.00523. PubMed PMID: 27288366; PubMed Central PMCID: PMCPMC4972278.

4. Li Z, Defoort J, Tasdighian S, Maere S, Van de Peer Y, De Smet R. Gene Duplicability of Core Genes Is Highly Consistent across All Angiosperms. Plant Cell. 2016;28(2):326–44. doi: 10.1105/tpc.15.00877. PubMed PMID: 26744215; PubMed Central PMCID: PMCPMC4790876.

5. Hua Z, Zou C, Shiu SH, Vierstra RD. Phylogenetic comparison of F-Box (FBX) gene superfamily within the plant kingdom reveals divergent evolutionary histories indicative of genomic drift. PLoS One. 2011;6(1):e16219. Epub 2011/02/08. doi: 10.1371/journal.pone.0016219. PubMed PMID: 21297981; PubMed Central PMCID: PMCPMC3030570.

6. Hua Z, Doroodian P, Vu W. Contrasting duplication patterns reflect functional diversities of ubiquitin and ubiquitin-like protein modifiers in plants. Plant J. 2018;95(2):296–311. Epub 2018/05/09. doi: 10.1111/tpj.13951. PubMed PMID: 29738099.

7. Force A, Lynch M, Pickett FB, Amores A, Yan YL, Postlethwait J. Preservation of duplicate genes by complementary, degenerative mutations. Genetics. 1999;151(4):1531–45. PubMed PMID: 10101175; PubMed Central PMCID: PMCPMC1460548.

8. He X, Zhang J. Rapid subfunctionalization accompanied by prolonged and substantial neofunctionalization in duplicate gene evolution. Genetics. 2005;169(2):1157–64. doi: 10.1534/genetics.104.037051. PubMed PMID: 15654095; PubMed Central PMCID: PMCPMC1449125.

9. Conant GC, Birchler JA, Pires JC. Dosage, duplication, and diploidization: clarifying the interplay of multiple models for duplicate gene evolution over time. Current Opinion in Plant Biology. 2014;19:91–8. doi: 10.1016/j.pbi.2014.05.008. PubMed PMID: 24907529.

10. Hua Z, Pool JE, Schmitz RJ, Schultz MD, Shiu SH, Ecker JR, et al. Epigenomic programming contributes to the genomic drift evolution of the F-Box protein superfamily in Arabidopsis. Proc Natl Acad Sci U S A. 2013;110(42):16927–32. Epub 2013/10/02. doi: 10.1073/pnas.1316009110. PubMed PMID: 24082131; PubMed Central PMCID: PMCPMC3801079.

11. McHale L, Tan X, Koehl P, Michelmore RW. Plant NBS-LRR proteins: adaptable guards. Genome Biol. 2006;7(4):212. Epub 2006/05/09. doi: gb-2006-7-4-212 [pii] 10.1186/gb-2006-7-4-212. PubMed PMID: 16677430; PubMed Central PMCID: PMC1557992.

12. Nei M, Gu X, Sitnikova T. Evolution by the birth-and-death process in multigene families of the vertebrate immune system. Proc Natl Acad Sci USA. 1997;94:7799–806.

13. Nei M, Niimura Y, Nozawa M. The evolution of animal chemosensory receptor gene repertoires: roles of chance and necessity. Nat Rev Genet. 2008;9(12):951–63. Epub 2008/11/13. doi: nrg2480 [pii] 10.1038/nrg2480. PubMed PMID: 19002141.

14. Shiu S-H, Shih M-C, Li W-H. Transcription factor families have much higher expansion rates in plants than in animals. Plant Physiol. 2005;139(1):18–26. PubMed PMID: 16166257.

15. Zou C, Lehti-Shiu MD, Thibaud-Nissen F, Prakash T, Buell CR, Shiu S-H. Evolutionary and expression signatures of pseudogenes in *Arabidopsis* and rice. Plant Physiol. 2009;151(1):3–15. PubMed PMID: 19641029.

16. Marone D, Russo MA, Laido G, De Leonardis AM, Mastrangelo AM. Plant nucleotide binding site-leucine-rich repeat (NBS-LRR) genes: active guardians in host defense responses. Int J Mol Sci. 2013;14(4):7302–26. doi: 10.3390/ijms14047302. PubMed PMID: 23549266; PubMed Central PMCID: PMCPMC3645687.

17. Burge CB, Karlin S. Finding the genes in genomic DNA. Curr Opin Struct Biol. 1998;8(3):346–54. Epub 1998/07/17. PubMed PMID: 9666331.

18. Mathe C, Sagot MF, Schiex T, Rouze P. Current methods of gene prediction, their strengths and weaknesses. Nucleic Acids Res. 2002;30(19):4103–17. Epub 2002/10/05. PubMed PMID: 12364589; PubMed Central PMCID: PMCPMC140543.

19. Coghlan A, Fiedler TJ, McKay SJ, Flicek P, Harris TW, Blasiar D, et al. nGASP--the nematode genome annotation assessment project. BMC Bioinformatics. 2008;9:549. Epub 2008/12/23. doi: 10.1186/1471-2105-9-549. PubMed PMID: 19099578; PubMed Central PMCID: PMCPMC2651883.

20. Guigo R, Flicek P, Abril JF, Reymond A, Lagarde J, Denoeud F, et al. EGASP: the human ENCODE Genome Annotation Assessment Project. Genome Biol. 2006;7 Suppl 1:S2 1–31. Epub 2006/08/24. doi: 10.1186/gb-2006-7-s1-s2. PubMed PMID: 16925836; PubMed Central PMCID: PMCPMC1810551.

21. Schweikert G, Zien A, Zeller G, Behr J, Dieterich C, Ong CS, et al. mGene: accurate SVM-based gene finding with an application to nematode genomes. Genome Res. 2009;19(11):2133–43. Epub 2009/07/01. doi: 10.1101/gr.090597.108. PubMed PMID: 19564452; PubMed Central PMCID: PMCPMC2775605.

22. Behr J, Bohnert R, zZeller G, Schweikert G, Hartmann L, Ratsch G. Next generation genome annotation with mGene.ngs. BMC Bioinformatics. 2010;11(Suppl 10):08.

23. van Dijk EL, Jaszczyszyn Y, Naquin D, Thermes C. The Third Revolution in Sequencing Technology. Trends Genet. 2018;34(9):666–81. Epub 2018/06/27. doi: 10.1016/j.tig.2018.05.008. PubMed PMID: 29941292.

24. Birney E, Clamp M, Durbin R. GeneWise and Genomewise. Genome Res. 2004;14(5):988–95. Epub 2004/05/05. doi: 10.1101/gr.1865504. PubMed PMID: 15123596; PubMed Central PMCID: PMCPMC479130.

25. Altschul SF, Gish W, Miller W, Myers EW, Lipman DJ. Basic local alignment search tool. Journal of Molecular Biology. 1990;215(3):403–10. doi: 10.1016/S0022-2836(05)80360-2. PubMed PMID: 2231712.

26. Li W, Godzik A. Cd-hit: a fast program for clustering and comparing large sets of protein or nucleotide sequences. Bioinformatics. 2006;22(13):1658–9. Epub 2006/05/30. doi: btl158 [pii] 10.1093/bioinformatics/btl158. PubMed PMID: 16731699.

27. Hua Z, Vierstra RD. The cullin-RING ubiquitin-protein ligases. Annu Rev Plant Biol. 2011;62:299–334. Epub 2011/03/05. doi: 10.1146/annurev-arplant-042809-112256. PubMed PMID: 21370976.

28. Gingerich DJ, Hanada K, Shiu SH, Vierstra RD. Large-scale, lineage-specific expansion of a bric-a-brac/tramtrack/broad complex ubiquitin-ligase gene family in rice. Plant Cell. 2007;19(8):2329–48. Epub 2007/08/28. doi: 10.1105/tpc.107.051300. PubMed PMID: 17720868; PubMed Central PMCID: PMCPMC2002615.

29. Kent WJ. BLAT-the BLAST-like alignment tool. Genome Res. 2002;12(4):656–64. Epub 2002/04/05. doi: 10.1101/gr.229202. Article published online before March 2002. PubMed PMID: 11932250; PubMed Central PMCID: PMC187518.

30. Lewin HA, Robinson GE, Kress WJ, Baker WJ, Coddington J, Crandall KA, et al. Earth BioGenome Project: Sequencing life for the future of life. Proc Natl Acad Sci U S A. 2018;115(17):4325–33. Epub 2018/04/25. doi: 10.1073/pnas.1720115115. PubMed PMID: 29686065; PubMed Central PMCID: PMCPMC5924910.

31. Yang X, Kalluri UC, Jawdy S, Gunter LE, Yin T, Tschaplinski TJ, et al. The F-box gene family is expanded in herbaceous annual plants relative to woody perennial plants. Plant Physiol. 2008;148(3):1189–200. Epub 2008/09/09. doi: 10.1104/pp.108.121921. PubMed PMID: 18775973; PubMed Central PMCID: PMCPMC2577272.

